# Rapid evolution and horizontal gene transfer in the genome of a male-killing *Wolbachia*

**DOI:** 10.1101/2020.11.16.385294

**Authors:** Tom Hill, Robert L. Unckless, Jessamyn I. Perlmutter

**Affiliations:** 4055 Haworth Hall, The Department of Molecular Biosciences, University of Kansas, 1200 Sunnyside Ave, Lawrence, KS, 66045

**Keywords:** *Wolbachia*, *Drosophila innubila*, male killing, genome evolution, phage WO

## Abstract

*Wolbachia* are widespread bacterial endosymbionts that infect a large proportion of insect species. While some strains of this bacteria do not cause observable host phenotypes, many strains of *Wolbachia* have some striking effects on their hosts. In some cases, these symbionts manipulate host reproduction to increase the fitness of infected, transmitting females. Here we examine the genome and population genomics of a male-killing *Wolbachia* strain, *w*Inn, that infects *Drosophila innubila* mushroom-feeding flies. We compared *w*Inn to other closely-related *Wolbachia* genomes to understand the evolutionary dynamics of specific genes. The *w*Inn genome is similar in overall gene content to *w*Mel, but also contains many unique genes and repetitive elements that indicate distinct gene transfers between *w*Inn and non-*Drosophila* hosts. We also find that genes in the *Wolbachia* prophage and Octomom regions are particularly rapidly evolving, including those putatively or empirically confirmed to be involved in host pathogenicity. Of the genes that rapidly evolve, many also show evidence of recent horizontal transfer among *Wolbachia* symbiont genomes, suggesting frequent movement of rapidly evolving regions among individuals. These dynamics of rapid evolution and horizontal gene transfer across the genomes of several *Wolbachia* strains and divergent host species may be important underlying factors in *Wolbachia*’s global success as a symbiont.

## Introduction

*Wolbachia* are the most widespread endosymbionts on the planet, infecting an estimated 40-52% of all insect species (Zug and Hammerstein 2012; Weinert *et al.* 2015). These obligate intracellular Gram-negative α-proteobacteria of the order *Rickettsiales* infect the gonads of their hosts and are primarily transmitted vertically via the cytoplasm from mother to offspring (Hertig and Wolbach 1924; Serbus and Sullivan 2007). *Wolbachia* of insects and other arthropods have adopted cunning techniques to facilitate their matrilineal spread by manipulating host reproduction to increase the proportion of infected, transmitting females in the population (Werren *et al.* 2008; Hurst and Frost 2015). The most common form of this reproductive parasitism is cytoplasmic incompatibility (CI), where crosses between infected males and uninfected females result in death of offspring. If the mother is also infected with a compatible strain, offspring are rescued from death, giving infected females a fitness advantage in the population over uninfected females (Yen and Barr 1971; Turelli and Hoffman 1991; Sinkins *et al.* 1995). Three other, less common forms of reproductive parasitism rely on sex ratio distortion to increase the proportion of transmitting females each generation. These phenotypes are known as parthenogenesis (asexual reproduction of females, (Russell and Stouthamer 2011)), feminization (genetic males physically develop and reproduce as females, (Bouchon *et al.* 1998; Kageyama *et al.* 2002)), and male killing (infected males die, (Hurst *et al.* 1999; Dyson *et al.* 2002)). Additional forms of transmission, including horizontal transfer between hosts of different species or strains (Vavre *et al.* 1999; Haine *et al.* 2005), are less understood and thought to be comparatively rare, but are likely key to *Wolbachia*’s ubiquitous spread around the world (Sanaei *et al.* 2020).

The incredible success of *Wolbachia* in becoming one of the world’s most widespread infections (Werren *et al.* 2008; LePage and Bordenstein 2013) is in part due to its diverse genetic toolkit. Indeed, *Wolbachia* strains are so diverse that they are divided into many supergroup clades, of which 18 have been named (Taylor *et al.* 2018; Laidoudi *et al.* 2020; Lefoulon *et al.* 2020). Studies on strains in supergroups A and B are the best represented in the literature, and include many reproductive parasite strains of hosts such as mosquitoes and a large number of *Drosophila* species (Gerth *et al.* 2014). Much of the diversity between *Wolbachia* genomes is contributed from prophage WO, the genome of phage WO of *Wolbachia* that has inserted itself into the bacterial chromosome and replicates along with core *Wolbachia* genes. Prophage WO sometimes retains the potential to form infective phage particles later, and sometimes degrades over time losing the potential to form new viral particles (Metcalf *et al.* 2014; Bordenstein and Bordenstein 2016). Prophages and phages are highly mobile and dynamic elements in the genome, often picking up new genes via horizontal gene transfer, and many such unique prophage WO genes have the potential to confer important functions in interactions with the eukaryotic host (Bordenstein and Bordenstein 2016). Indeed, functional and evolutionary analyses of the genetic loci that underlie CI have shown that they are in fact prophage WO genes that interact directly with the eukaryotic host to manipulate reproductive and developmental processes (Beckmann *et al.* 2017; LePage *et al.* 2017; Lindsey *et al.* 2018; Beckmann *et al.* 2019). Related to phage WO is a cassette of 8 genes known as Octomom. This cassette contains paralogs of phage WO genes but replicates separately and copy number of Octomom regions is correlated with regulation of *Wolbachia* titer, making strains more or less pathogenic to the host (Chrostek and Teixeira 2015; Duarte *et al.* 2020).

Despite the great diversity and interest in a variety of *Wolbachia* infections, most research attention has focused on CI, largely due to its use in vector control strategies around the globe (Zabalou *et al.* 2004). These programs take advantage of the natural abilities of *Wolbachia* to both block viral transmission and spread itself via reproductive parasitism (Hedges *et al.* 2008; Teixeira *et al.* 2008; O’neill *et al.* 2018; Mains *et al.* 2019; Ross *et al.* 2019). Comparatively fewer analyses have been done on *Wolbachia* genomes of strains that induce male killing (Dyer and Jaenike 2004; Duplouy *et al.* 2013; Metcalf *et al.* 2014). However, male killing merits additional analysis due to its potential in vector control (Berec *et al.* 2016), role in shaping arthropod evolution (Jiggins *et al.* 2000), and the close relationship between CI and male killing (Dyer *et al.* 2005). Indeed, the CI genetic loci are located only a few genes away from the male-killing candidate gene, *wmk* (*WO-mediated killing*) in the *Wolbachia* strain of *Drosophila melanogaster* (*w*Mel) (Perlmutter *et al.* 2019). Also, many male-killing and CI strains are closely related (Sheeley and Mcallister 2009), and several strains are multipotent in that they can switch between the two phenotypes either within the same host or between different hosts (Hurst *et al.* 2000; Sasaki *et al.* 2002; Jaenike 2007). The close genetic relationship between the two phenotypes indicates that studies on male killing may inform CI and vice versa. In addition, their overall similarities may help narrow down evolutionary dynamics that are unique to each phenotype or shared between them.

Among the few *Wolbachia* male-killers of flies is the strain infecting *Drosophila innubila, w*Inn (Dyer and Jaenike 2004). This strain is particularly interesting as it is closely related to *Wolbachia* found in the main *Drosophila* model species, *w*Mel of *D. melanogaster*, which causes CI (Sheeley and Mcallister 2009). In addition, the symbiosis between *w*Inn and its host has been maintained for thousands of years, and despite this, there is no evidence of host resistance to the phenotype in modern populations (Jaenike and Dyer 2008; Unckless and Jaenike 2012). Due to the close relationship between *w*Mel and *w*Inn and the longstanding symbiosis of *w*Inn with its host, analysis of *w*Inn population dynamics can be used to uncover evolutionary trends that may be important in reproductive parasitism generally, male-killers specifically, or other interactions with the host.

Here, we sequence the genome of the *Wolbachia* strain infecting *D. innubila*, *w*Inn, and conduct population genomic analyses using sequences from 48 *Wolbachia-*infected individual wild females from four populations in Arizona. We demonstrate overall similarity of the genome content with *w*Mel, but with several dozen unique genes implying horizontal gene transfer with divergent hosts. We determine if genes from prophage and Octomom regions, including those thought to be involved in various host-microbe interactions, show more evidence of adaptive evolution than background genes, consistent with *Wolbachia*’s ability to rapidly adapt to diverse hosts. Finally, we examine population structure and co-inheritance of *Wolbachia* with mitochondria to determine if horizontal transmission occurs frequently in *w*Inn.

## Methods

### Genome sequence of *w*Inn

For a single *Wolbachia*-positive strain described previously (Unckless and Jaenike 2012), we extracted DNA following the protocol described in (Chakraborty *et al.* 2017). We prepared the DNA as a sequencing library using the Oxford Nanopore Technologies Rapid 48-h (SQK-RAD002) protocol, which was then sequenced using a MinION (Oxford Nanopore Technologies, Oxford, UK; NCBI SRA: TBD) (Jain *et al.* 2016). The same DNA was also used to construct a fragment library with insert sizes of ∼180bp, and we sequenced this library on an Illumina HiSeq 4000 (150 bp paired-end, Illumina, San Diego, CA, NCBI SRA: TBD).

Oxford Nanopore sequencing read bases were called post hoc using the built in read_fast5_basecaller.exe program with options: –f FLO-MIN106 –k SQK-RAD002 –r–t 4. We assembled the raw Oxford Nanopore sequencing reads alongside the Illumina paired-end short sequencing reads using SPAdes version 3.13.0 (Bankevich *et al.* 2012), which generated an initial assembly of 83 contigs. We then attempted to improve this initial assembly using the 83 assembled fragments, along with Nanopore sequencing reads and Illumina paired-end short sequencing reads in MaSuRCA version 3.4.1 (Zimin *et al.* 2013), defining the expected genome size as 1.5 million bp. This produced a single contig 1,286,799 bp long. We then used Pilon version 1.23 to polish the genome with minion fragments for 3 iterations (Walker *et al.* 2014) and further polished with Racon version 1.4.3 for three iterations using the short read data (Vaser *et al.* 2017). We then verified the contiguity of the assembly using BUSCO version 3.0 (SIMÃO *et al.* 2015). From a search for 221 proteobacteria orthologs, we found 181 complete single copy orthologs and 2 fragmented orthologs (compared to 180 complete and 2 fragmented for the published *w*Mel genome: NC_002978.6).

### Fly collections and *Wolbachia* infection confirmation

We collected wild *Drosophila* at four mountainous locations across Arizona between the 22nd of August and the 11th of September 2017 (Hill and Unckless 2020b; Hill and Unckless 2020a). Specifically, we collected at the Southwest research station in the Chiricahua mountains (31.871 latitude, −109.237 longitude), Prescott National Forest (34.540 latitude, −112.469 longitude), Madera Canyon in the Santa Rita mountains (31.729 latitude, −110.881 longitude) and Miller Peak in the Huachuca mountains (31.632 latitude, −110.340 longitude). Baits consisted of store-bought white button mushrooms (*Agaricus bisporus*) placed in large piles about 30cm in diameter, at least 5 baits per location. A sweep net was used to collect flies over the baits in either the early morning or late afternoon between one and three days after the bait was left. Flies were sorted by sex and our best guess of species based on morphology at the University of Arizona and were flash frozen at −80°C before being shipped on dry ice to Lawrence, KS. Specifically, we separated individuals likely to be *Drosophila innubila* from the rest of the collections for further processing and genetic confirmation of species identification.

We further analyzed the 343 putative *D. innubila* flies which we homogenized and extracted DNA from using the Qiagen Gentra Puregene Tissue kit (USA Qiagen Inc., Germantown, MD, USA) (Hill and Unckless 2020b; Hill and Unckless 2020a). We tested these samples for infection using *Wolbachia* primers specific to the *Wolbachia surface protein* (*wsp*) gene alongside a positive and negative control (Zhou *et al.* 1998).

The reaction mixture for the *wsp* PCR consisted of 1uL DNA, 1u 10X buffer (Solis Biodyne), 1.0 μl of 20 mM MgCl2 (Solis Biodyne), 1 μl of dNTPs (20 μM each), 0.5 μl of forward (F) primer (81F 5’-TGGTCCAATAAGTGATGAAGAAAC-3’, 20 μM), 0.5 μl of reverse (R) primer (691R 5’-AAAAATTAAACGCTACTCCA-3’, 20 μM), 0.5 μl of Taq DNA polymerase (5 U/μl) (Solis Biodyne) and water to make up the final volume of 10 μl. The amplification reaction consisted of one cycle of 1 min at 94°C, 1 min at 58°C and 2 min at 72°C, followed by 35 cycles of 15 s at 94°C, 1 min at 58°C and 2 min at 72°C, and one cycle of 15 s at 94°C, 1 min at 58°C and 7 min at 72°C. These conditions yielded 610 basepair (bp) PCR products, which we observed running out the product on a 1% agarose TAE gel. This survey yielded 48 *Wolbachia*-positive lines.

For the 48 *Wolbachia*-infected lines we previously extracted DNA and sequenced the host and *Wolbachia* genomes on two runs of an Illumina HiSeq 4000 (150 bp paired-end (Hill and Unckless 2020a; Hill and Unckless 2020b), Illumina, San Diego, CA), producing an average of 20,618,752 reads per sample, of which an average of 436,527 mapped to *Wolbachia* per sample, as summarized in Supplementary Table 1.

### Genome annotation

We annotated the *w*Inn genome using Prokka version 1.15.4 (Seemann 2014), detecting 2686 total genes, of which 1390 were retained following size and quality filtering (> 50bp, quality score > 20, Supplementary Table 2). Using this annotation of the genome, we extracted coding sequences and generated amino acid sequences using GFFread version 0.12.1 (Pertea 2011). We also downloaded the coding sequence and amino acid sequences for open reading frames in the *Wolbachia* of *Drosophila melanogaster* (Canton S strain) (*w*Mel-CS, SAMN02604000), the *Wolbachia* of *Drosophila simulans* (Riverside strain) (*w*Ri, SAMN02603205), the *Wolbachia* of *Drosophila simulans* (Hawaii strain) (*w*Ha, SAMN02604273), and the *Wolbachia* of *Culex pipiens* (*w*Pip, SAMN02296948). We used blastp version 2.9.0 (Altschul *et al.* 1990) to identify orthologs for these genes in *w*Inn (parameters: hsp = 1, num_alignments = 1, e-value < 0.00001). For each set of orthologs we created a gene alignment using MAFFT version 7.409 (parameters: --auto) and for 100 randomly chosen genes made a visual inspection of amino acid sequences to confirm similarity of putative orthologous sequences. We then verified the completeness of the extracted amino acids sequences using BUSCO version 3.0 (SIMÃO *et al.* 2015). From a search for 221 proteobacteria orthologs, we found 185 complete single copy orthologs and 1 fragmented ortholog, and 1 complete and duplicated ortholog (compared to 184 complete, 1 duplicated and 2 fragmented for the published *w*Mel genome: NC_002978.6).

To annotate the repetitive content of the *w*Inn genome, we used RepeatModeler version 2.0.1 (Smit and Hubley 2008) and RepeatMasker version 4.0.9 (parameters: -gff -gccalc -s -norna) (Smit and Hubley 2013-2015).

### Genomic variation in *wInn*

For all 48 *Wolbachia*-positive lines collected in 2017, we mapped short reads to the *D. innubila* genome (Hill *et al.* 2019), masked using RepeatMasker version 4.0.9 (parameters: -gff -gccalc -pa 4 -s) (Smit and Hubley 2013-2015), a custom library of *D. innubila* repeats (Hill *et al.* 2019), and the masked *w*Inn genome using BWA MEM version 0.7.17-r1188 (Li and Durbin 2009) and SAMtools version 1.9 (Li *et al.* 2009). We then extracted aligned reads mapping to *w*Inn and used GATK version 4.0.0 to remove optical and PCR duplicates and realign around indels (Mckenna *et al.* 2010; Depristo *et al.* 2011). We then called variants in the *w*Inn genome of each *Wolbachia*-positive lines using GATK HaplotypeCaller version 4.0.0 (Mckenna *et al.* 2010; Depristo *et al.* 2011), considering only variants with a quality score greater than 500. Finally, we combined VCFs using BCFtools version 1.7 (Narasimhan *et al.* 2016) to create a multiple sample VCF.

### Detection of selection on *Wolbachia* genes

For each *w*Inn gene with an ortholog in *w*Ha and *w*Ri, we generated an alignment of the coding sequence of each gene using MAFFT version 7.409 (parameters: --auto). Following this alignment, we reformatted the alignment into a PAML version 1.3.1 usable format and generated a gene tree using PRANK version 0.170427 (parameters: +F -showtree -d=paml) (LÖytynoja 2014). We next used codeML (Yang 2007) to calculate the non-synonymous divergence (dN) and synonymous divergence (dS) across the entire gene tree and find the best fitting branches model (Model 7 or 8), as well as calculate dN and dS on each branch of the tree (Model 1), specifically looking at the estimates of dN/dS on the *w*Inn branch versus all other branches. For both the total tree and specifically the wInn branch, we looked for gene functional categories with higher dN/dS than all other genes, after controlling for gene length.

### Population structure across *w*Inn populations

For synonymous sites in the VCF, we used VCFtools version 0.1.16 (Danecek *et al.* 2011) to calculate the fixation index (Fst) between each population and the other populations (Brown 1970). We also performed a principle component analysis on the variation found across the samples in R version 3.5.1 (Team 2013), using the VCF input as a presence/absence matrix.

### Ancient and recent horizontal transfer

We reasoned that if no horizontal gene transfer was occurring, then *Wolbachia* variation would be perfectly linked to mitochondrial variation, while non-vertical transfer would break that pattern. To assess this, we looked at all pairwise combinations of mitochondrial and *Wolbachia* alleles and recorded loci with all four allele sets across the two loci across the 48 samples (e.g. GT, AT, GC and AC), giving a recombination like signature (suggesting non-vertical inheritance). We then counted the number of discordant and non-discordant SNPs in 10-kbp windows across the *w*Inn genome to identify specific sections enriched for discordant SNPs. We used a χ^2^ test to identify specific functional categories enriched for discordant SNPs.

For long term horizontal gene transfer, we used the VHICA R package version 0.2.7 to calculate dS vs codon bias for all pairwise for all shared genes for pairwise combinations of *w*Inn, *w*Ha and *w*Ri (Wallau *et al.* 2016). We reasoned that horizontal transfer of a gene from a highly divergent *Wolbachia* would produce a signal of increased dS between *w*Ha and *w*Inn for that gene and could polarize which species had the horizontal transfer event based on the dS of that gene in the pairs *w*Inn-*w*Ha, *w*Ha-*w*Ri and *w*Inn-*w*Ri. We considered dS to be excessively high in a gene if it was greater than the mean dS + the variance for that window of effective number of codons (5 codons window size, sliding 5 codons) (Wallau *et al.* 2016). We considered a gene to be a putative horizontal acquisition in *w*Inn if dS compared to *wHa* and *w*Ri is excessively high compared, but dS is also not significantly higher when comparing *w*Ha to *w*Ri. We then performed a χ^2^ test to look for functional categories that are enriched for putatively horizontally acquired genes.

Finally, we assessed the extent of ancient horizontal transfer across the *Wolbachia* phylogeny. We downloaded all *Wolbachia* genomes and their annotations from the NCBI genome database (summarized in Supplementary Table 1), based on the known NCBI annotations we found groups of orthologous genes. We generated codon alignments for these orthologous genes using MAFFT (parameters: --auto) (Katoh *et al.* 2002), and generated a gene tree for each gene using PhyML (model = GTR, gamma = 8, bootstraps = 100) (Guindon *et al.* 2010). We also generated a whole species phylogeny for these genomes and to place *w*Inn on the phylogeny. For all genes found in all species with high confidence (231 genes), we generated a multigene phylogeny with 100 bootstraps using PhyML (model = GTR, gamma = 8, bootstraps = 100). We then used CADM.global in APE (Paradis *et al.* 2004) to assess the extent of species/gene tree discordance to test for consistency between phylogenies, with the null hypothesis that the phylogenies are different across 100,000 permutations per species/gene tree comparison (so a significant *p*-value will suggest little discordance between phylogenies). Finally, we performed a χ^2^ test to look for functional categories that are enriched for putatively horizontally transferred genes.

## Results

### *w*Inn genome assembly reveals a genome similar to *w*Mel and evidence of horizontal gene transfer from multiple host genera

*D. innubila* is a mycophagous species in the *Drosophila* subgenus, found throughout the southwestern USA and northwestern Mexico on mountain-top forests known as ‘Sky Islands’, separated by large expanses of desert (Jaenike *et al.* 2003; Dyer and Jaenike 2005; Dyer *et al.* 2005; Jaenike and Dyer 2008). Since the *Wolbachia* strain of *Drosophila innubila* (*w*Inn) is one of the few *Wolbachia* strains known to cause male killing in *Drosophila* (Jaenike et al. 2003; Dyer and Jaenike 2005), studying its evolutionary and population dynamics allows new genetic insights into male-killing populations. To that end, we examined the genome and population genomic variation in *w*Inn. In a previous survey we collected wild *D. innubila* from four geographically isolated mountain locations and tested strains for *Wolbachia* using PCR to amplify *wsp* primers and found 48 lines infected with *Wolbachia* (Supplementary Table 1, 13 from the Chiricahua mountains, 27 from Prescott, 2 from the Huachucas, and 6 from the Santa Ritas) (Hill and Unckless 2020b; Hill and Unckless 2020a).

We sequenced and assembled the genome using a combination of short and long reads for one strain. The *w*Inn genome is a single chromosome 1,247,635 base pairs long, with 35.1% GC content (Figure 1A). We found 1390 genes, 1331 found previously in other *Wolbachia*: 1292 genes are shared with *w*Mel, 1248 shared with *w*Rec, and 1059 shared with *w*Ri. We found the *w*Inn genome had a BUSCO score of 81.9% (181 complete single copy orthologs and 2 fragmented orthologs, from a sample of 221) compared to 81.3% in *w*Mel (180 complete single copy orthologs and 2 fragmented orthologs, from a sample of 221). Of the 1331 previously identified genes, 954 are conserved across all four genomes (Figure 1C, Supplementary Table 2), including 12 prophage WO-A and 54 prophage WO-B genes in all genomes (Supplementary Table 3), and 9 Octomom genes (5 orthologs to *w*Mel and 4 paralogs of these, genes linked to *Wolbachia* pathogenicity) (Chrostek and Teixeira 2015). Interestingly, these Octomom genes are not found in a single cassette like in *w*Mel but are instead spread throughout the genome (Figure 1A). The genes orthologous to WO-B genes are found in 3 groups (Figure 1A, called WOInn-B1, WOInn-B2 and WOInn-B3). Despite the fragmentation of the prophage regions, the genes are syntenic to the prophage-B region in *w*Mel. We also found 10 Type IV secretion pump genes, found in two cassettes, as in *w*Mel (Figure 1A). Consistent with previous results, *w*Inn is closely related to *w*Mel within supergroup A, clustering with other supergroup A *Wolbachia* genomes (Figure 1B, Supplementary Figure 1).

**Figure 1.**
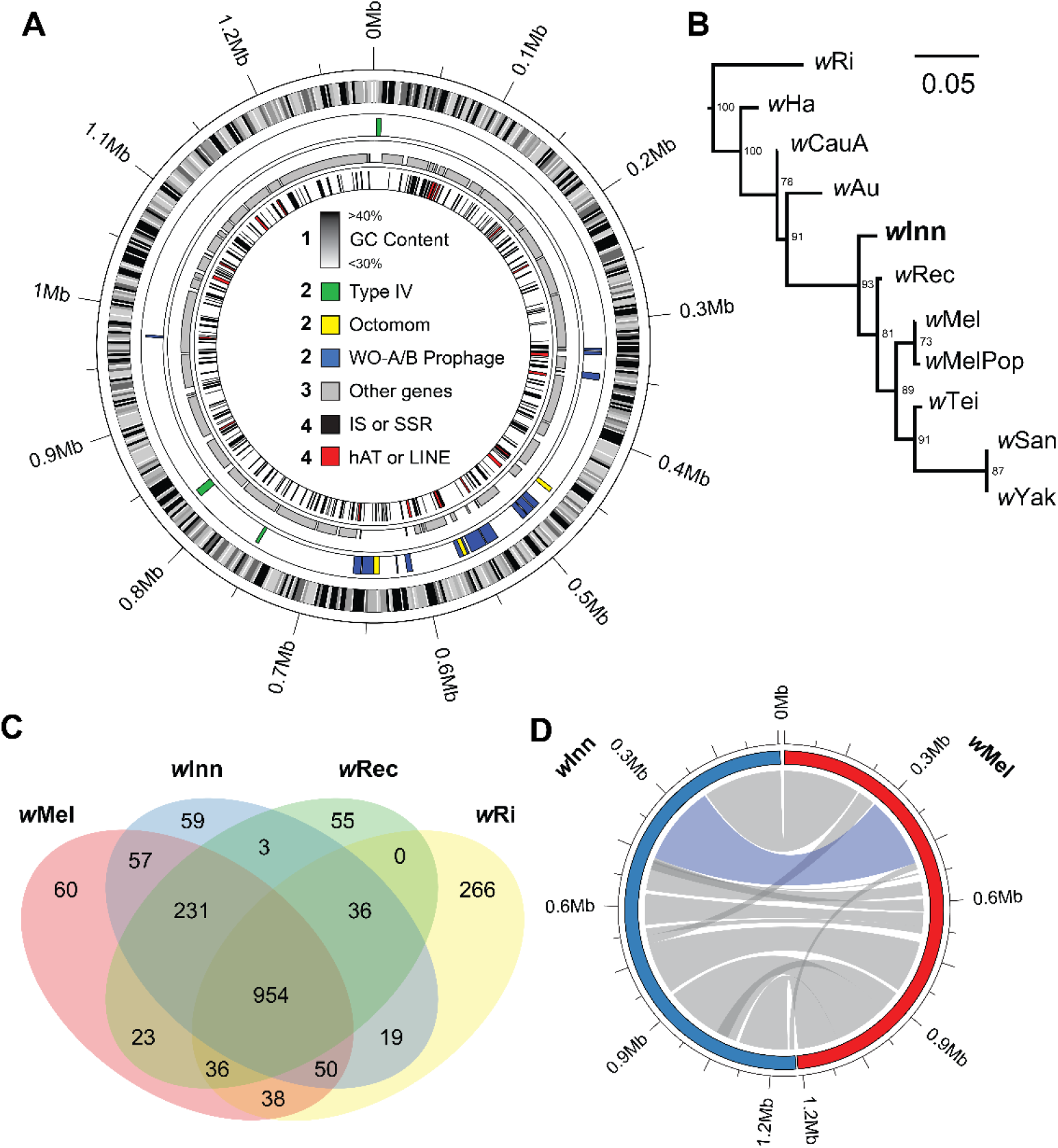
**A.** Genome schematic of the *w*Inn genome. Circles correspond to the following: 1) GC content of the *w*Inn genome in 10-kbp windows, between 30% and 40%. Darker colors have higher GC content. 2) Locations of genes thought to interact with hosts, specifically prophage orthologous to WO-A and WO-B genes in *w*Mel (blue), Type IV secretion pumps (green), and Octomom genes (yellow). 3) Loci of non-phage genes. 4) Loci of repetitive content, with short simple repeats and interspersed satellites (SSR and IS, black), and hAT or LINE TE insertions (red). **B.** Phylogeny of *Wolbachia* genomes closely related to *w*Inn for reference, as a subset of Supplementary Figure 1. Bootstrap support of each branch is shown on the nodes (of 100 bootstraps). A description of each *Wolbachia* genome and the species they infect is given in Supplementary Table 1. **C.** The overlap of genes between *w*Inn, *w*Mel, and *w*Ri. **D.** Synteny between *w*Mel and *w*Inn, with single large inversion shown in blue, while consistent synteny groups are shown in grey.

We found 3 genes are shared between the male-killing *w*Inn genome and *w*Rec (which reportedly kills males when introgressed into a sister species, but causes CI in its native host (Jaenike 2007)), but absent in the *w*Mel and *w*Ri genomes, both of which induce CI in their native hosts (Figure 1C). All three genes are hypothetical proteins found in other non-male-killing *Wolbachia* supergroup A genomes (including other varieties of *w*Mel). These genes are therefore not likely to be involved with male-killing specifically, but have been gained in the ancestral supergroup A, then lost in *w*Mel-CS and *w*Ri genomes. *w*Inn does not appear to have a reduced, relic prophage genome like *w*Rec, and instead shares most prophage genes with *w*Mel despite the more distant phylogenetic relationship (Figure 1C, (Metcalf *et al.* 2014)). The 57 genes absent in *w*Rec but present in *w*Inn and *w*Mel consists of 21 prophage genes, 4 transcription genes, 9 metabolism genes, and 21 genes of unknown function. We attempted to further confirm the differences in genomic content by mapping short reads from *w*Rec, *w*Mel, *w*Inn and *w*Ha to each of the genomes pairwise and find the exact same number of genes shared in each case, supporting the assembled genomes are not missing any shared genes. The *w*Rec genome appears to be missing portions of the regions orthologous to 0.35-0.55Mb and 1.24-1.28Mb in the *w*Inn genome, which also includes a large portion of the prophage WO-B genome.

Of the 59 coding sequences in *w*Inn, but not found in other *Wolbachia*, 33 of these have high similarity to mRNA in *Formica* wood ants that may have an overlapping range with *D. innubila* (non-redundant megablast e-value < 0.00005) (Francoeur 1973; Altschul *et al.* 1990). *D. innubila* also contains multiple transposable element (TE) sequences shared with *Camponotus* (a genus within *Formica*) (Hill *et al.* 2019), and so horizontal transfer may occur frequently between these species. Mites are a potential vector for horizontal transfer of genes and *Wolbachia* (Houck *et al.* 1993; Brown and Lloyd 2015) and in keeping with this, we find 7 new *w*Inn genes have a high similarity to *Varroa destructor* transcriptome sequences (non-redundant megablast e-value < 0.00005, Supplementary Table 3). Though this is not a common *Drosophila* mite, it may be closely related to other *Drosophila* mites without sequenced genomes that may vector genes between *Wolbachia* strains. Alternatively, it may be a source of other *Wolbachia* infections in *D. innubila*, or horizontal exchange of genes between *Wolbachia* strains (Brown and Lloyd 2015). These genes could also be undescribed mobile elements which have spread from a different species, like the mobile elements described below. The nine remaining sequences have no known orthologs. Among the 157 genes absent in *w*Inn but present in *w*Mel, we found no functional categories enriched (*p*-value > 0.12).

We found 12.38% of the *w*Inn genome is repetitive, similar to other *Wolbachia* (Figure 1A) (Foster *et al.* 2005; Woolfit *et al.* 2013); most of these sequences are short simple repeats, satellites, and insertions from 16 bacterial insertion sequences (selfish elements found in bacteria). 1.49% of the genome consists of insertions of a single hAT family element (*hobo*-like DNA transposon found in *Drosophila*, rnd-1_family-6) inserted in 14 loci across the genome and 3.74% consists of 3 LINE elements inserted in 37 loci across the genome (long interspersed nuclear elements, an RNA transposon order found in *Drosophila*), primarily in clusters (Figure 1A) (Wicker *et al.* 2007). Consistent with the potential horizontal transfer seen for several genes, we find one LINE element (rnd-1_family-12 with 14 insertions) is homologous to a LINE found previously in *Varroa destructor* (or a close relative), while another (rnd-1_family-165 with 12 insertions) is homologous to a LINE found in *Formica* wood ants (non-redundant megablast e-value < 0.00005) (Altschul *et al.* 1990). No homologous sequence can be identified for the hAT element (non-redundant megablast e-value = 1 no hits), which is also the only TE found as complete sequences suggesting a more recent horizontal acquisition of full active copies, while most of the LINE element insertions are degraded, supporting ancient horizontal acquisitions (Supplementary Figure 2).

### Octomom and prophage genes are rapidly evolving in *w*Inn and other *Wolbachia*

We next used PAML to determine genes rapidly evolving in *w*Inn compared to the closely related *Wolbachia* genomes (Yang 2007). For each ortholog set, we identified the proportion of synonymous (dS) substitutions and amino acid changing, nonsynonymous substitutions (dN) (per possible synonymous or nonsynonymous substitution, respectively) occurring on each branch of the phylogeny to identify changes between the gene sequence of *w*Inn, *w*Ha, and *w*Ri. We expect dN/dS to be higher when genes are faster evolving, due to more nonsynonymous fixations, potentially due to positive selection (Yang 2007). We chose *w*Ha and *w*Ri over *w*Mel or other genomes as these genomes are diverged enough from *w*Inn to provide some signal (unlike *w*Mel or *w*Rec, where dS ~ 0.001 between genomes), while not being diverged enough to have too little similarity or saturated rates of dS. Genes previously suspected to be involved in pathogenicity (certain Octomom and prophage genes), are rapidly evolving on all branches (Figure 2A, GLM t-value = 2.750, *p*-value = 0.0061), while DNA metabolism genes are more rapidly evolving exclusively on the *w*Inn branch, though not significantly (Figure 2A, GLM *t-*value = 1.868, *p*-value = 0.0622). The fastest evolving DNA metabolism genes in *w*Inn are the non-phage genes WD1095 (*radC*, a DNA repair protein), WD0065 (a DNA binding protein), WD0057 (a host integration factor) and WD0752 (*xerC*, a recombinase).

**Figure 2.**
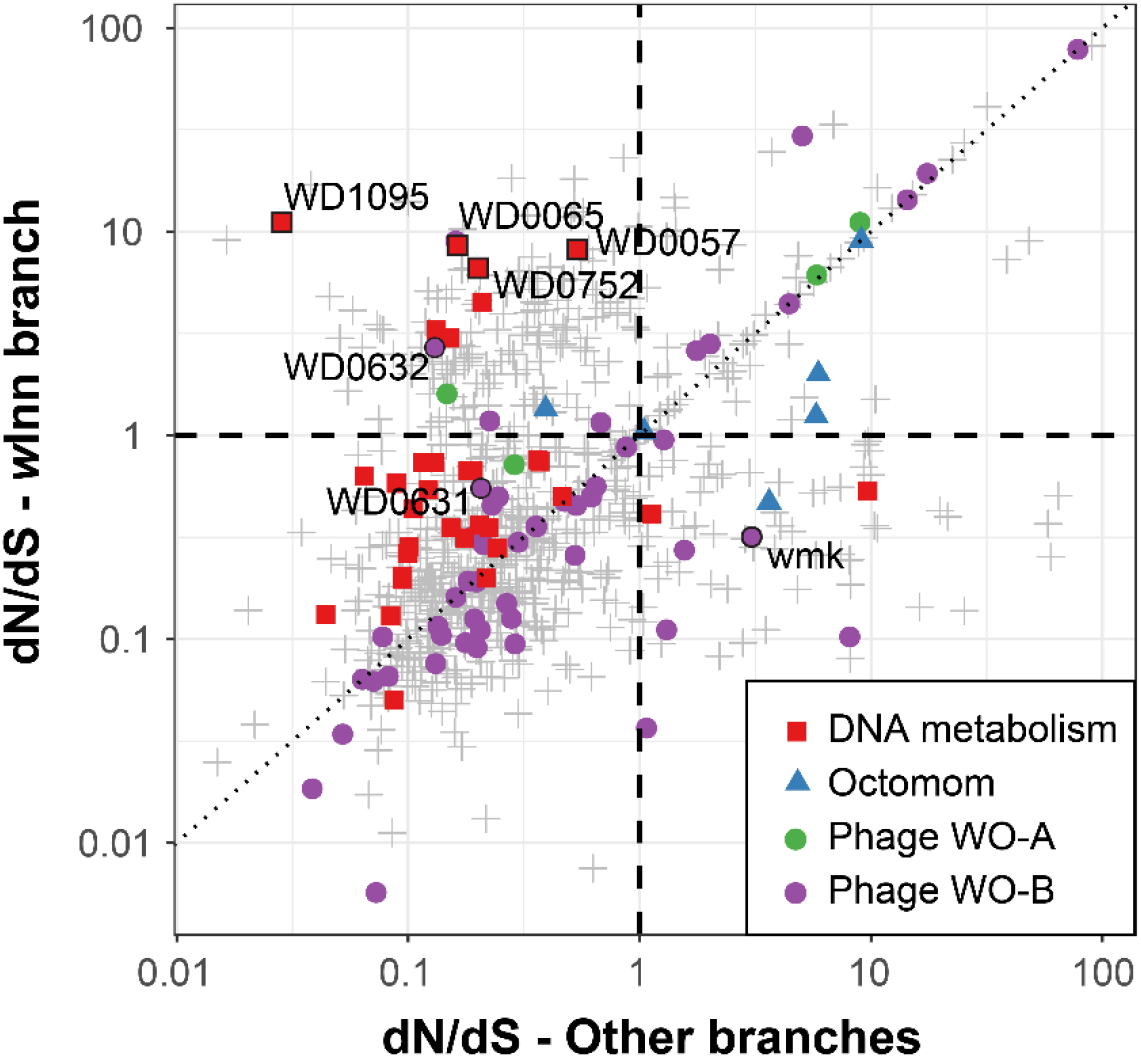
Rate of evolution of the *w*Inn branch versus evolution on the *w*Ri/*w*Ha branches. Functional categories of interest (DNA metabolism genes, prophage genes and Octomom genes) are highlighted by different shapes and colors. Dashed lines show dN/dS = 1 for both axes, while the dotted line shows where dN/dS is equal on the axes. Genes of interest, either due to putative involvement in *Wolbachia* pathogenicity, or due to high dN/dS in *w*Inn exclusively are named and labelled with a black outline to distinguish them.

Several genes have been previously implicated in reproductive parasitism in *Wolbachia* in *Drosophila*, so we specifically examined the evolution of these genes in *w*Inn and the other genomes (*wmk*: WD0626, *cifA*: WD0631, *cifB*: WD0632). These genes are evolving faster than background rates in *w*Inn (dN/dS = 0.27-1.56 in *w*Inn, versus dN/dS background median = 0.225), though not significantly (GLM t-value = 0.224, *p*-value = 0.642). Additionally, *wmk* is only rapidly evolving on the *w*Ha and *w*Ri branches (dN/dS = 1.56), while *cifB* (a gene thought to be involved with cytoplasmic incompatibility) (LePage *et al.* 2017), is only rapidly evolving in *w*Inn (Figure 2, WD0632, dN/dS = 1.10). We also examined the rate of evolution of specific codons in these genes but find no specific sites are driving this rapid evolution in these putative host manipulation genes (*p*-value > 0.05). The Type IV secretion genes are also faster evolving than the background across the total phylogeny, but not significantly (GLM t-value = 1.427, *p*-value = 0.154).

### Some prophage genes show evidence of recent horizontal transfer in *w*Inn and across the *Wolbachia* phylogeny

Horizontal transfer is frequently occurring in microbes at different levels: symbiotic bacteria such as *Wolbachia* can switch hosts to propagate in new species and can switch between individual hosts within a species (Vavre *et al.* 1999; Haine *et al.* 2005; Riegler *et al.* 2005; Werren *et al.* 2008; Ilinsky 2013). Beyond this, specific genes and regions of genomes can horizontally transfer to and from bacteria of the same strain allowing for a recombination-like process which may facilitate adaptation, or can transfer to different strains which allows for the acquisition of new genes, allowing for adaptation to better propagate within their hosts, in a process known as horizontal gene transfer (Lawrence 1999; Dutta and Pan 2002).

We suspected that many symbiont genes potentially involved in unique *w*Inn host-microbe interactions may be more likely to horizontally transfer between *Wolbachia* strains than other genes. So, we next looked for evidence of horizontal gene transfer since the divergence of *w*Inn from *w*Ha and *w*Ri. We used VHICA (Wallau *et al.* 2016) to compare synonymous divergence (dS) to the effective number of codons for the pairwise comparisons of *w*Inn-*w*Ha, *w*Inn-*w*Ri, and *w*Inn-*w*Ha for genes with orthologs in all three species. As dS is constrained by codon usage bias, it can be higher when the effective number of codons is higher, and so should be controlled for when comparing the divergence of two different genes (Wallau *et al.* 2016). If a gene has horizontally transferred into *w*Inn from another *Wolbachia* (but not *w*Ha or *w*Ri), we expect the dS to be higher in these comparisons than expected, after controlling for codon usage bias (Wallau *et al.* 2016). i.e. there will be elevated dS in both comparisons involving *w*Inn, but not the *w*Ha-*w*Ri comparison. We found 13 genes have elevated dS in just the *w*Inn comparisons (Figure 3, Supplementary Table 4), suggesting potential horizontal gene acquisition from another *Wolbachia*. These genes are enriched for prophage WO-A & WO-B genes (χ^2^ test, χ^2^ = 60.476, df = 1, *p*-value = 7.448e-15), and Octomom genes (χ^2^ test, χ^2^ = 181.64, df = 1, *p*-value = 2.2e-16) that have elevated divergence in *w*Inn. The 6 genes not associated with the prophage or Octomom regions are all genes of unknown function. We did not suspect horizontal transfer of the *Wolbachia* organism to play a role in the elevated divergence seen here due to the similarity in dS for the *w*Inn-*w*Ri and *w*Ha-*w*Ri comparisons (Wilcoxon Rank Sum Test W = 462970, *p*-value = 0.7687).

**Figure 3.**
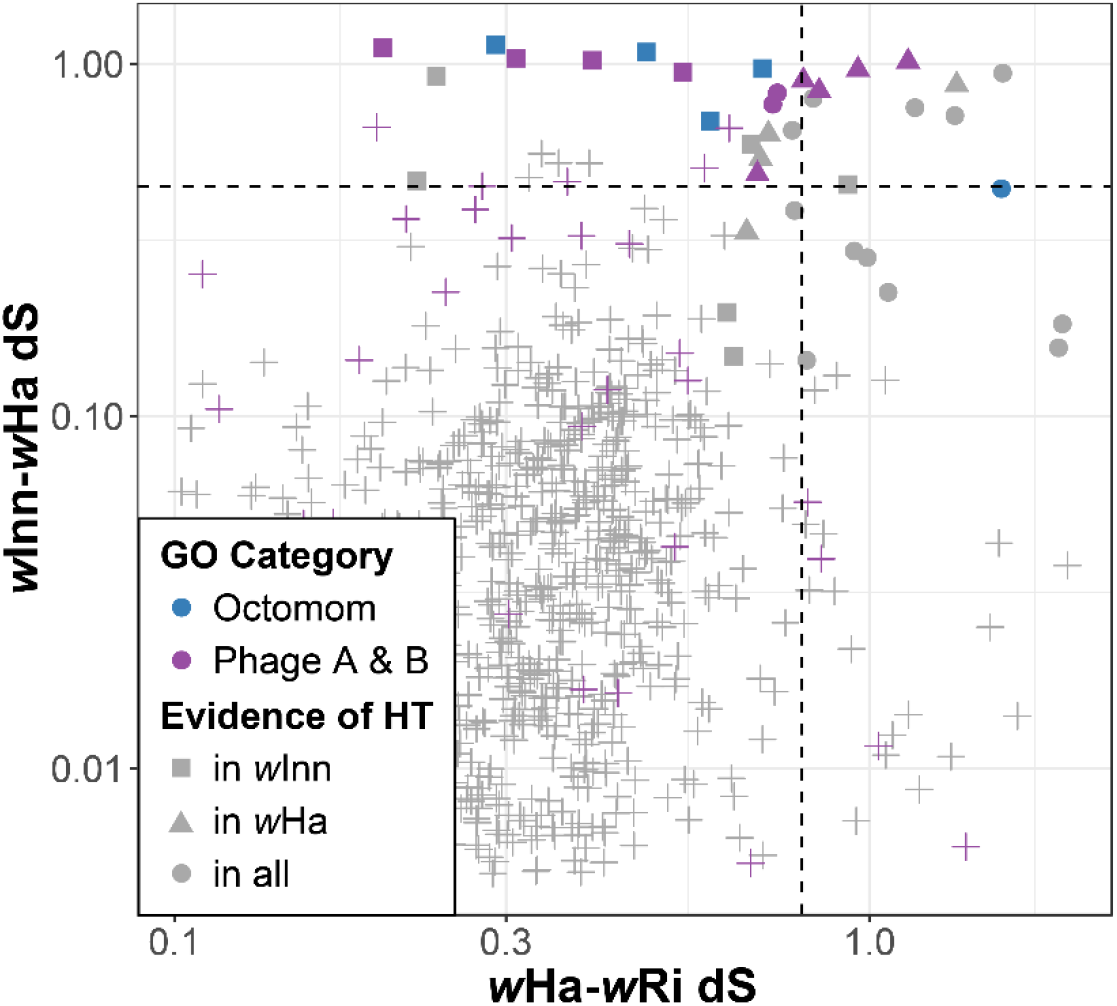
Comparison of dS between pairs of *Wolbachia* suggesting horizontal transfer of genes. Point colors show the gene ontology categories (GO category) for Octomom genes and Prophage WO-B genes. Point shape indicates evidence of excessive divergence (and possible horizontal transfer) in either *w*Inn, *w*Ha, or both. Dashed lines show the mean + the variance for each axis as a vague cutoff for elevated synonymous divergence. Note this does not consider effective number of codons, so it is not the cutoff used to determine if there is elevated divergence (which is difficult to illustrate as it differs per gene), but it is a close approximation.

To examine if these gene categories frequently horizontally transfer, or if these transfers are unique to *w*Inn, we downloaded 54 *Wolbachia* genomes (all genomes available for download on NCBI genomes, described in Supplementary Table 1) and made gene alignments for all orthologs and attempted to identify gene tree/species tree discordance. We assumed that excessive gene tree/species tree discordance would be due to large amounts of horizontal gene transfer. We attempted to look for functional categories which show more tree discordance than expected and across 847 orthologous genes, and found excessive amounts of discordance for prophage WO genes (Table 1, 36 of 47 genes, degrees of freedom = 1, χ^2^ = 111.1, *p*-value = 5.62e-26) and Octomom genes (Table 1, 7 of 7 genes, degrees of freedom = 1, χ^2^ = 71.27, *p*-value = 3.395e-17) across large evolutionary distances, while no other categories have significantly more discordance than expected. We find a significant overlap in the genes which have horizontally transferred in *w*Inn and across the whole phylogeny (χ^2^ = 49.003, *p*-value = 2.556e-12), suggesting that specific (prophage) genes are more likely to horizontally transfer than others and demonstrating that the recent horizontal transfers are not unique to this strain.

**Table 1:** Summary of species tree/gene tree discordance analysis. Using 847 orthologous genes across 54 genomes (Supplementary Table 5), we identified genes which showed significant discordance from the species tree. Table shows the number of genes in each category showing discordance, and if this discordance is significant using a χ^2^ test, using an expected number of discordant genes per category based on the size of the category.

| Functional Category | Discordant genes | Genes in category | $\chi^2$ | $p$-value |
| --- | --- | --- | --- | --- |
| Biosynthesis | 1 | 50 | 2.374 | 0.876 |
| Cell envelope | 1 | 31 | 0.952 | 0.671 |
| Cellular processes | 1 | 35 | 1.238 | 0.732 |
| DNA metabolism | 1 | 42 | 1.759 | 0.815 |
| Energy metabolism | 0 | 90 | 7.438 | 0.006 |
| Octomom | 7 | 7 | 71.278 | 3.395e-17 |
| Phage WO-A | 12 | 13 | 111.105 | 5.626e-26 |
| Phage WO-B | 24 | 33 | 165.927 | 5.817e-38 |
| Protein synthesis | 1 | 144 | 9.985 | 0.002 |
| Regulatory functions | 0 | 11 | 0.909 | 0.340 |
| Transcription | 0 | 52 | 4.298 | 0.038 |
| Transport and binding proteins | 0 | 44 | 3.636 | 0.057 |
| Type IV | 0 | 10 | 0.826 | 0.363 |
| Unknown function | 22 | 285 | 0.102 | 0.749 |

### *w*Inn is highly structured between locations, and is not always co-inherited with the mitochondria

We next mapped the short-read data for 48 samples of wild *D. innubila* infected with *w*Inn, collected from four locations, to the repeat masked *w*Inn genome and called polymorphism. From this, we identified 30 SNPs as coding synonymous, 69 SNPs as coding non-synonymous, and 235 SNPs as non-coding across all individuals. The *w*Inn samples are highly structured based on both the total and synonymous variation (Figure 4). Using a principle component analysis, we find three clear clusters, separating the Chiricahua and Prescott populations, and grouping the Santa Rita and the Huachuca populations together (Figure 4), as seen with the mitochondrial genome and consistent with previous findings (Jaenike *et al.* 2003; Dyer and Jaenike 2005; Jaenike and Dyer 2008; Hill and Unckless 2020a). This suggests that females are not moving between locations or at least not reproducing after moving, creating structure in the populations of this maternally inherited endosymbiont (Jaenike *et al.* 2003).

**Figure 4:**
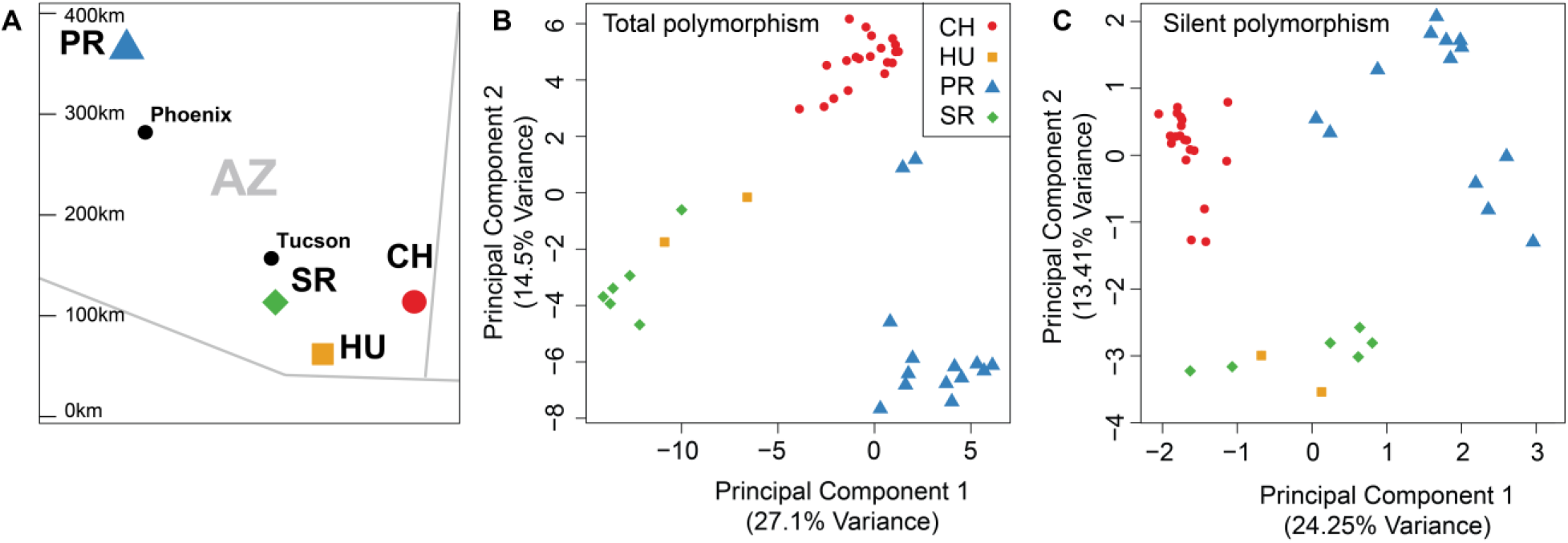
Principal component analysis of genetic variation in *w*Inn **A.** Map of locations samples were taken from in this survey, adapted from (Hill and Unckless 2020a). Phoenix and Tucson are shown as points of reference. **B.** total polymorphism and **C.** silent polymorphism in *w*Inn samples, colored and shaped by location of collection, Chiricahuas (CH, red circles), Huachucas (HU, orange squares), Prescott (PR, blue triangles) and Santa Ritas (SR, green diamonds).

However, when building a maximum-likelihood tree of the *w*Inn samples using all polymorphisms in the core *Wolbachia* genes, we find some evidence of migration between populations (Supplementary Figure 3A). Specifically, we find two samples from PR cluster within CH and share the CH mitochondrial haplotype, suggesting there may be multiple *Wolbachia* types shared between population locations due to host migration (Supplementary Figure 3). These 2 PR samples are also closer to CH than other PR samples in the principal component analysis (Figure 4). This signature is not seen in the host, likely due to the recent establishment of *D. innubila* (Hill and Unckless 2020a), particularly in Prescott.

To identify if specific factors are contributing to the population structure, we calculated the fixation index (F_ST_), a measure of pairwise divergence between a subpopulation and the total population, between the three clustered groups. We expect FST to be elevated in cases where SNPs are found at high frequencies in a single population but not the remaining samples. As expected with the non-recombining bacterial genome, we found signatures of FST are uniform genome wide, with no specific windows of elevated FST compared to the rest of the genome (1kbp windows, GLM *p*-value > 0.432) and no functional categories are enriched for high or low FST (Supplementary Table 6, GLM *p*-value > 0.611).

When comparing the inheritance of the maternally transmitted *w*Inn and *D. innubila* mitochondria, we found little evidence of discordant inheritance, consistent with a previous study (Dyer and Jaenike 2005). We do however find evidence of three clusters of 49 SNPs in the *w*Inn genome which show evidence of recombination-like events (all four allele combinations between the *Wolbachia* site and the mitochondria site, Supplementary Figure 4), suggesting either imperfect co-inheritance of the *w*Inn and mitochondria, recurrent mutation, or a horizontal transfer event. The two larger clusters are contained within the prophage WO-A and WO-B portions of the genome, suggesting horizontal movement of the prophage is the cause of this discordance. We also find the copy number of the prophage regions differs between *w*Inn lines: the prophage region has significantly higher sequence coverage in 8 of 48 lines, varying from 1-4x the average coverage of the *Wolbachia* genome (Wilcoxon Rank Sum W > 443221, *p*-value < 0.04, Figure 5). In addition, phage copy number is negatively correlated with *Wolbachia* titer. Indeed, higher phage coverage correlates with lower *Wolbachia* titer up to a point, consistent with potentially active phage lysing symbiont cells. This additional sequencing coverage of phage regions and negative correlation with titer despite a potentially weaker signal from not controlling for host age or other factors may suggest that the prophage region produces active phage particles. Active phage would also be consistent with horizontal transfer of prophage genes between hosts, as discussed above (Figure 5).

**Figure 5:**
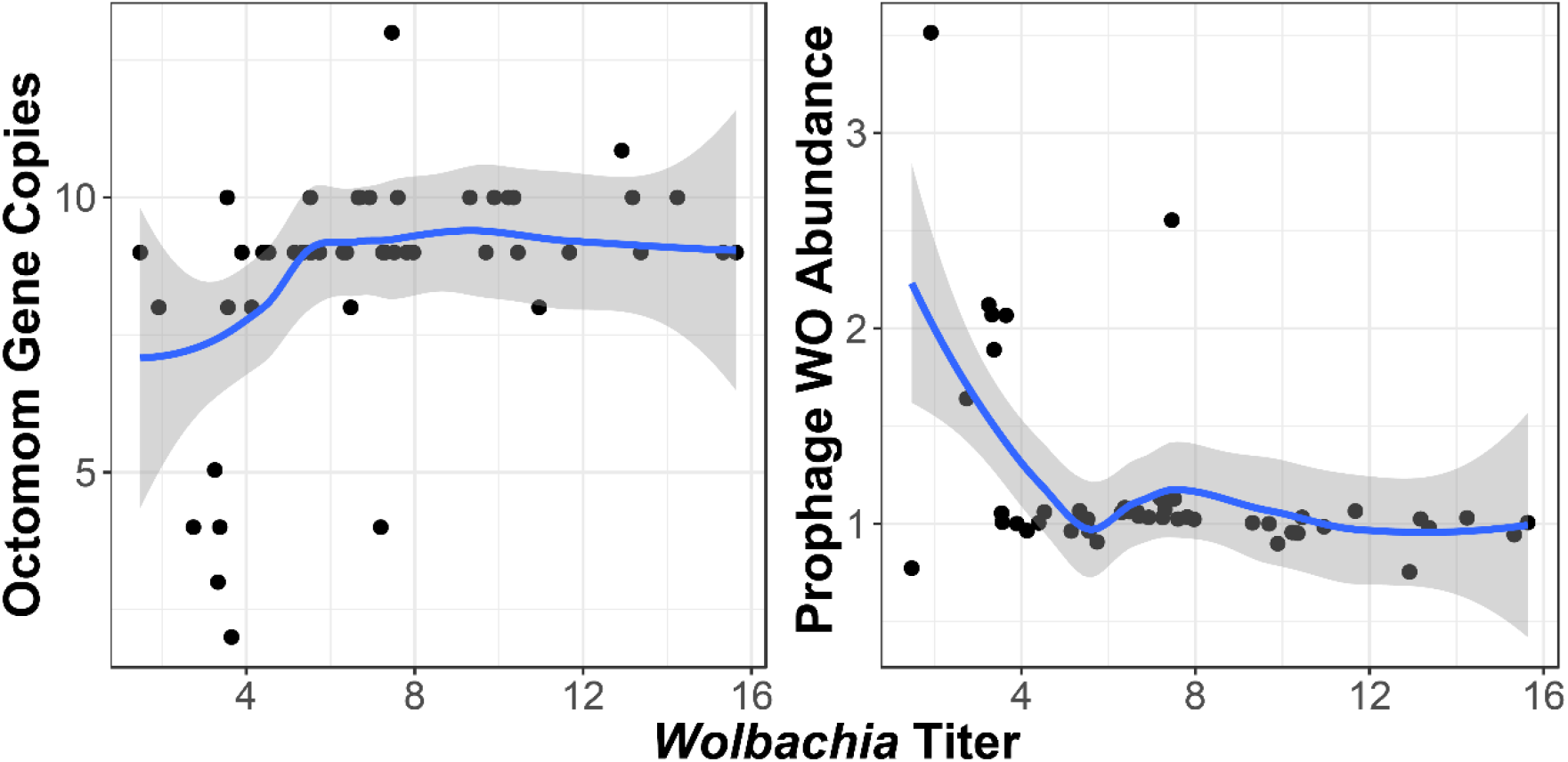
Octomom copies compared to *Wolbachia* titer (total *Wolbachia* coverage/host autosomal coverage) and prophage abundance genes (prophage coverage/total *Wolbachia* coverage) compared to the *Wolbachia* titer (total *Wolbachia* coverage/host autosomal coverage).

Given the rapid evolution of Octomom genes, and the previously identified association between increased pathogenicity and Octomom copy number, we also examined the effect of copy number on *Wolbachia* titer within lines. We found all strains contain Octomom genes in a cassette, though coverage of the Octomom genes appears to also differ between lines, consistent with potentially differing copy numbers of Octomom (Supplementary Figure 5). We also identified a positive correlation between Octomom copy number and *Wolbachia* titer, though it is not significant (Supplementary Figure 5, GLM t-value = 1.930, *p*-value = 0.0597). However, *Wolbachia* titer does not seem to be associated with Octomom copy number once Octomom copy number is greater than 9 (Supplementary Figure 5, GLM t-value = 0.534, *p*-value = 0.596). This suggests that while putatively increased Octomom copy number may be important for titer, it has a reduced effect higher than ~8 copies. The differences in titer are not significantly associated with any SNPs or gene duplications (*p*-value > 0.186).

## Discussion

We sequenced and assembled the genome of the *Wolbachia* strain infecting *Drosophila innubila*, *w*Inn. *w*Inn is one of the few strains of *Wolbachia* to cause male-killing in *Drosophila* (Jaenike *et al.* 2003) and so we sought to examine the evolutionary dynamics of its genome, with particular focus on prophage and Octomom regions that have genes putatively or empirically involved in *Wolbachia* pathogenicity (Metcalf *et al.* 2014; Chrostek and Teixeira 2015; Bordenstein and Bordenstein 2016; Beckmann *et al.* 2017; LePage *et al.* 2017; Perlmutter *et al.* 2019). The genome content and dynamics of *w*Inn are largely like other closely-related strains (Figure 1), sharing most of its genome with similar supergroup A *Wolbachia* strains. *w*Rec is relatively closely related to *w*Inn and has been reported to kill males when introgressed into a sister host species, *D. subquinaria* (Jaenike 2007). When comparing the genomes of *w*Inn and *w*Rec to the other closely related strains in this analysis (all CI-inducing), we identify only 3 unique genes. However, these genes are found in other *Wolbachia* strains not used in our initial analyses, based on the non-redundant BLAST database (Altschul *et al.* 1990; Pruitt *et al.* 2005). The lack of unique male-killer genes is consistent with the idea that male killing is often hidden. Indeed, many strains like *w*Rec do not cause male killing in their native hosts but do kill males when transferred to other hosts or vice versa (Fujii *et al.* 2001; Sasaki *et al.* 2002; Jaenike 2007; Hughes and Rasgon 2014). In addition, there is evidence of host resistance that suppresses the phenotype in many systems, where the arms race between host and bacteria leads to loss of phenotype (Hornett *et al.* 2006; Jaenike 2007; Majerus and Majerus 2010). These factors indicate that absence of phenotype does not necessarily correlate with absence of symbiont genotype and male killing is instead also heavily dependent on host background (i.e. male killing is not a simple matter of presence/absence of a male-killing toxin gene). The fact that the *wmk* male-killing gene candidate is found in many non-male killer genomes also supports the idea that male-killers do not necessarily have unique genetic content involved in the phenotype and a combination of host- and microbe-dependent factors are necessary for the phenotype to occur (Perlmutter *et al.* 2019).

We also find that while the overall genetic content of *w*Inn is similar to *w*Mel, it has key differences compared to the more similar *w*Rec strain. The *w*Inn and *w*Mel strains share 57 total genes that are unique to them among those analyzed (including *w*Rec), of which 21 were prophage genes. These differences are most likely due to the loss of phage regions in *w*Rec (Metcalf *et al.* 2014). It is intriguing to note that *w*Rec contains relic phage regions that have lost many genes compared to those in *w*Mel and *w*Inn, and is likely unable to produce viral particles because of the absence of key virus structural genes (Metcalf *et al.* 2014). Meanwhile, *w*Inn and *w*Mel have not (Figure 1). Both *w*Inn and *w*Rec are closely related, both are capable of male killing (Jaenike *et al.* 2003; Metcalf *et al.* 2014), and both infect mycophagous species, yet one has an eroded prophage region while the other does not (Figure 1). In addition, evidence here based on higher sequence coverage and the inverse relationship between phage titer and *Wolbachia* titer, further suggests that there may be active phage WO particle production (Supplementary Figure 5). It is unclear why *w*Inn and *w*Mel would putatively maintain viable phage particle production while *w*Rec would not. Future research will be needed to determine any functional consequences of phage particles that may be playing a role in their retainment or loss across different systems.

We also identified 59 genes unique to the *w*Inn genome, and most intriguingly, 33 of these are homologous to *Formica* wood ant genes, an additional 7 share homology with *Varroa destructor* mites, and there are several TEs with homology to those in *Camponotus* ants. Along with evidence of horizontal gene transfer and rapid evolution, this homology with divergent hosts would suggest new possibilities for the genetic transfer routes in this system. Indeed, mites are known to transfer *Wolbachia* infections among *Drosophila* populations (Brown and Lloyd 2015), and *Varroa destructor* is a common parasitic species found throughout the United States and the rest of the world (Rosenkranz *et al.* 2010). Therefore, it is possible that this or a similar species has vectored either the entire *Wolbachia* symbiont or some genes among various arthropods, contributing to horizontal gene transfer in this strain. *Formica* wood ants and *Camponotus* ants are also common across the United States (Bolton *et al.* 2006), indicating that there is a possibility of interaction with the mites and/or *D. innubila*. The homology of *w*Inn genes with ant genes may indicate that there has been an exchange of genes or symbionts among these hosts, possibly via mites, and the ants and mites in the area may contain similar strains. In this model, the mites would vector either genes or symbionts among interacting hosts (Houck *et al.* 1993; Brown and Lloyd 2015), and the active phage particles could also play a role by primarily or excessively moving the prophage region among hosts, which may be much easier and more common than symbiont exchange.

Horizontal transfer of genes between *Wolbachia* strains and hosts is in line with existing literature demonstrating *Wolbachia*’s proclivity for genetic exchange. Indeed, the horizontal transfer of individual genes or the entire phage region among *Wolbachia* strains that is supported here reflects previously reported cases (Wang *et al.* 2016; Cooper *et al.* 2019). Phage WO genes appear to horizontally transfer between the *Wolbachia* genomes analyzed here more often than the rest of the genome, possibly due to the activity of the phage particles (Table 1, Figure 3), or the activity of surrounding mobile elements, as may have happened in the *D. yakuba* clade with horizontal transfer of the CI loci and nearby transposons (Cooper *et al.* 2019). Further, entire symbiont transfer may also potentially occur in this system, although it is rare if it does occur. Vertically transmitted symbionts, such as *Wolbachia*, are subject to strong selection within the host, and unlike frequently horizontally transferring microorganisms, can experience various degrees of genome reduction and other genetic adaptations that lead to essential ties with a specific host (Moran 2002; Dyer and Jaenike 2005; Jaenike and Dyer 2008; Mccutcheon and Moran 2012). However, bypassing this phenomenon, there are *Wolbachia* strains that can transfer to new hosts via various modes of transmission and then sweep rapidly across a new and sometimes divergent host population (Riegler *et al.* 2005; Baldo *et al.* 2008; Turelli *et al.* 2018; Sanaei *et al.* 2020). Whole symbiont transmission could be aided by frequent horizontal transfer of genes or entire regions, such as the prophage, just as we report here in *w*Inn. More broadly, acquisition of new genetic variants that the eukaryotic host is unfamiliar with may confer an advantage that could allow the *Wolbachia* to be transferred to a new host or maintained in an existing host. Indeed, some known cases of horizontal phage WO gene transfer among symbionts have functional effects on the symbiont’s ability to parasitize the host (Wang *et al.* 2016; Cooper *et al.* 2019). Most crucially, horizontal gene transfer in *Wolbachia* is not restricted to exchange among phages or bacteria, but also with the eukaryotic host. Phage WO stands unique among other described phages with its regions of large genes containing eukaryotic-like domains that imply both lateral transfer between animal and phage WO genomes and potential interaction with the eukaryotic host (Bordenstein and Bordenstein 2016). Among these genes in the prophage region known as the ‘Eukaryotic Association Module’ are the two CI loci and the male-killing gene candidate that empirically impact host reproductive biology (Beckmann *et al.* 2017; LePage *et al.* 2017; Perlmutter *et al.* 2019). Thus, the acquisition of ant and mite genes in *w*Inn reflects *Wolbachia*’s unique genetic exchanges more broadly as well as its tripartite interactions among phages, bacteria, and animals. Elucidating the function and fitness impacts of these genes, if any, will be an interesting area of future research. Also, if there is indeed frequent horizontal exchange in this system, this may then be a driving force for maintenance of phage particles, as they may confer selective advantages over time. Further supporting the idea of recurrent genetic exchanges in the *w*Inn ecosystem is evidence of both recently acquired TEs and more ancient, degraded TEs, some of which are homologous to those in *Formica* and *Varroa*.

Notably, we find a surprising amount of this repeat content in the *w*Inn genome, like other *Wolbachia* (Figure 1A, Supplementary Table 2). This is in contrast to other obligate intracellular parasites have relatively small, repeat free genomes (Woolfit *et al.* 2013). Most of the elements found in the *w*Inn genome are also shared with the host, *D. innubila* (Hill *et al.* 2019), suggesting that inefficient selection has not allowed these TEs to be maintained for extensive periods of time, but instead these elements are recent acquisitions (Yoshiyama *et al.* 2001), possibly vectored by prophage transmission. It is possible that eventually these TEs will be shed from the *w*Inn genome and similar repeat families have been acquired in the past and eventually gone extinct and removed from the genome, in a cyclical fashion (Maruyama and Hartl 1991; Lohe *et al.* 1995). Previous work in *w*MelPop suggests that reduced selection allowed repeats to accumulate in the *Wolbachia* genomes (Woolfit *et al.* 2013). This could also be the case for *w*Inn, allowing these families to be maintained in the genome for extended periods of time, as opposed to removed immediately. The lack of similarity between *w*Inn and *w*MelPop repeats suggests that mobile elements have not been maintained since the common ancestor of these two *Wolbachia* strains, and that turnover is much more frequent.

Beyond just genome content, we analyzed the evolutionary dynamics of the genes in *w*Inn. The overall finding, in line with what is known about phage biology, is that prophage and Octomom genes are indeed rapidly evolving along all *Wolbachia* branches analyzed, although not significantly so. There was also a trend of potentially more rapid evolution of DNA metabolism genes in *w*Inn (Figure 2). Specifically, the genes with the strongest signal were involved in DNA repair, DNA binding, host integration, and recombination (Supplementary Table 3). This could relate to what may be an unusual amount of horizontal gene transfer or high number of repetitive elements in this system, as DNA metabolism genes would aid in these dynamics. Looking more specifically at the *cifA*/*B* CI genes and the *wmk* male-killing gene candidate, we find that they individually also show evidence of rapid evolution across the phylogeny, although they are not more rapidly evolving than the rest of the prophage region (Figure 2). These are putatively or empirically demonstrated key genes in parasitizing host reproduction and may be in an adaptive arms race with the host (Beckmann *et al.* 2017; LePage *et al.* 2017; Lindsey *et al.* 2018; Perlmutter *et al.* 2019), an idea that is supported by the rapid evolution.

Regarding the Octomom regions, we find that these genes are rapidly evolving across all the *Wolbachia* genomes in the clade examined, as expected, not just in the male-killing *Wolbachia* (Figure 2). The rapid evolution of these genes may indicate they are frequently involved in host-pathogen interactions as suggested elsewhere (Chrostek and Teixeira 2015), and as is seen with immune genes and other genes involved in interspecies arms races (Dawkins and Krebs 1979; Sackton *et al.* 2007; Obbard *et al.* 2009; Palmer *et al.* 2018). Indeed, previous studies have found an association between Octomom copy number and increased titer (Chrostek and Teixeira 2015), as we see here (Supplementary Figure 5). Additionally, while prophage may be able to excise themselves for transfer, Octomom may utilize transposable elements for horizontal transfer (Chrostek and Teixeira 2015). In line with this, the Octomom genes are fragmented across the genome rather than remaining in full cassettes, and the horizontal transfer of transposable elements is more and more frequently being found to occur between species (Peccoud *et al.* 2017; Hill and Betancourt 2018; Wallau *et al.* 2018). Given the presence of *Varroa* repeat elements in the *w*Inn genome, this also provides a suitable vector organism to transfer these genes between *Wolbachia* types. When looking for evidence of recent horizontal transfer to *w*Inn more generally (after divergence from *w*Ha and *w*Ri), 13 genes were identified as potentially horizontally transferred to *w*Inn. Of these, 7 were prophage or Octomom genes (Figure 3), supporting all findings so far indicating particularly rapid evolution and frequent transfer of prophage and Octomom genes.

Finally, we compared the inheritance of mitochondria and *w*Inn in *D. innubila* populations, and find there is overall geographic structure in both cases (Figure 4) (Hill and Unckless 2020b; Hill and Unckless 2020a). However, we also find some discordance in inheritance in the *Wolbachia* prophage region, likely due to the activity of the prophage (Figure 5) resulting in extensive horizontal gene transfer of this region (Figure 4, Table 1, Supplementary Figure 4). When examining the phylogeny of *w*Inn genomes, we find some Prescott lines are grouped within Chiricahua lines, potentially driven by this horizontal gene transfer. Alternatively, since the populations are recently established (Hill and Unckless 2020b; Hill and Unckless 2020a), it may simply indicate that these lines and mitotypes differentially segregated into these populations upon establishment and that the types dominant in Chiricahuas are rarer in Prescott.

## Conclusion

Here, we provide the first genome description of *w*Inn of *D. innubila* and use population genomic analyses to understand the evolutionary dynamics of this symbiont. We show evidence of rapid evolution, particularly among prophage and Octomom regions. We also demonstrate that this system is potentially experiencing high rates of horizontal transfer of genes, phages, or entire symbionts. This may occur not only within or between *D. innubila* populations, but across divergent mite and ant species as well. These dynamics may contribute to the success of *Wolbachia* symbionts in these populations and may reflect a broader strategy for survival and adaptation in diverse *Wolbachia* around the globe.

## Acknowledgements

This manuscript was conceived and written thanks to suggestions and feedback from the Unckless lab at the University of Kansas. Collections were completed with assistance from Todd Schlenke at the University of Arizona, and the Southwest Research Station. The initial *w*Inn sample used for genome sequencing was graciously provided by John Jaenike. We thank Brittny Smith and Jenny Hackett at the KU CMADP Genome Sequencing Core (NIH Grant P20 GM103638) and K-INBRE Bioinformatics Core for assistance in genome isolation, library preparation, sequencing and computational resources.

## Data Availability

All sequencing data used in this study is available on the NCBI SRA under the project accession: PRJNA524688. Additional data regarding *D. innubila* population genomics is available in the following FigShare folder: https://figshare.com/account/home#/projects/87662 and the following DataDryad submission: https://datadryad.org/stash/share/wvfmDL39pdYrVUcgDFAfI33BOJu3KCJWuJyj-0M-qgA.

## Author contributions

TH devised analyses, performed genome assembly, performed analyses and wrote the manuscript, JIP devised analyses and wrote the manuscript, RLU devised analyses and wrote the manuscript.

## Ethics approval and consent to participate

Not applicable.

## Consent for publication

Not applicable.

## Supplementary Tables and Figures

**Supplementary Table 1:** Data (genomes and next generation sequencing information) used in this study. When applicable the number of sequences/reads and the proportion of these that are *Wolbachia* are shown in the table. SRA accession numbers are included for both short reads and genomes.

**Supplementary Table 2:** Genes annotated in the *w*Inn genome in GFF format, with their ID and *w*Mel ortholog when applicable. GFF also includes locations of transposable element insertions and short simple repeats.

**Supplementary Table 3:** dN/dS estimations between *w*Inn, *w*Ha and *w*Ri. Functional categories of genes are also noted in the table. Table includes both the non-synonymous (dN) and synonymous (dS) estimations across the total phylogeny of *w*Inn-*w*Ha-*w*Ri and on each branch of the phylogeny.

**Supplementary Table 4:** Table of VHICA output and divergence estimates. For each gene (using the *w*Inn gene name), we include the measure of effective codon usage (CUB), the dS pairwise between *w*Ha, *w*Inn and *w*Ri. If the gene has elevated dS, we note this. Table also include the gene ontology category of the gene.

**Supplementary Table 5:** Gene orthologs identified in each non-*w*Inn genome analysed, when looking for discordance from the whole species phylogeny. Gene annotation for each genome is given for each column. When the ortholog is absent from a genome it is labelled as NA.

**Supplementary Table 6:** FST enrichments for functional categories across the *w*Inn genome using a generalized linear model.

**Supplementary Figure 1.**
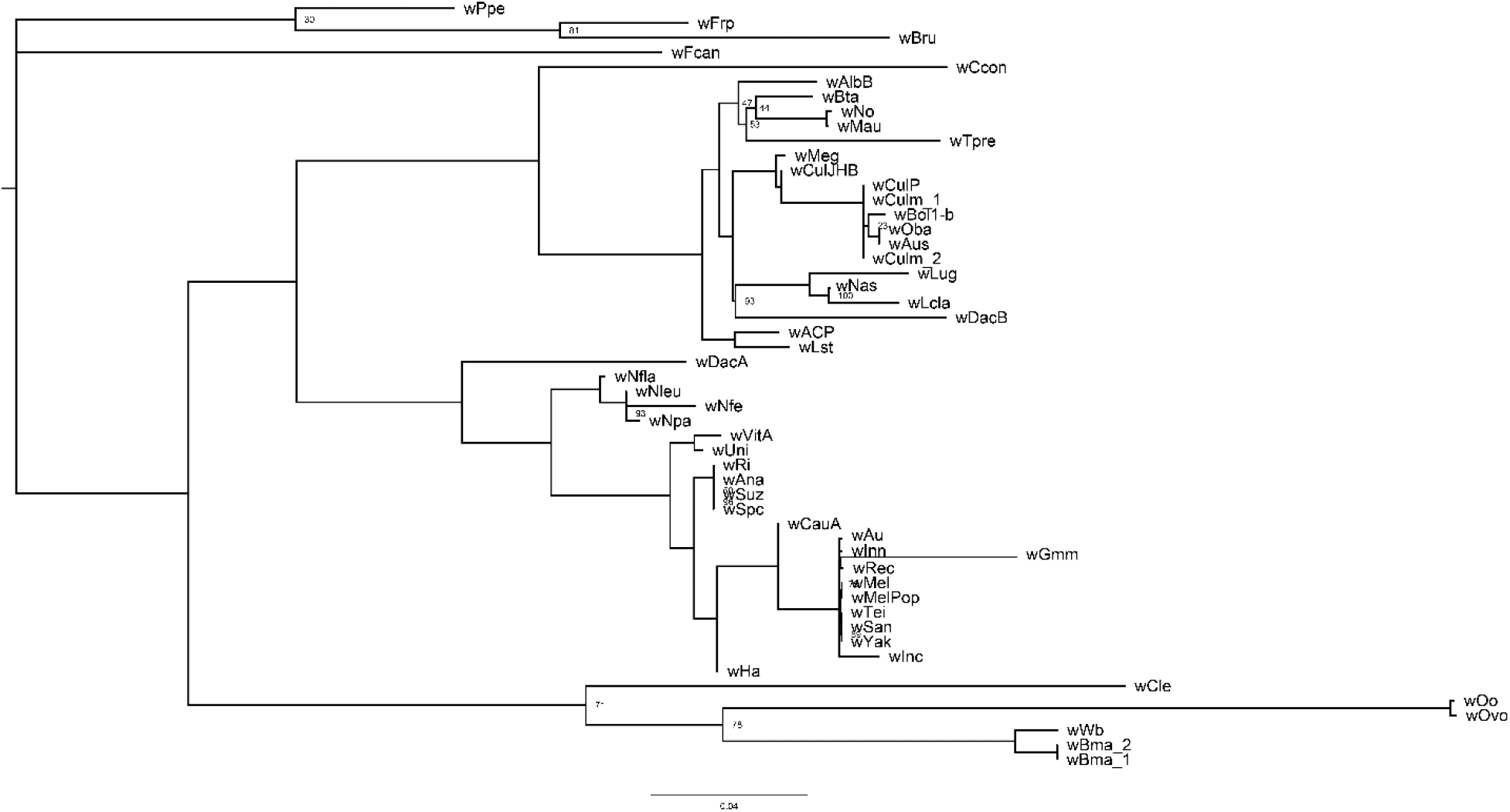
Phylogeny of *Wolbachia* genomes used in this survey. A subset of this phylogeny is used in Figure 1B. Nodes with 100 bootstrap support are not labelled, nodes with fewer than 100 bootstraps support are labelled.

**Supplementary Figure 2.**
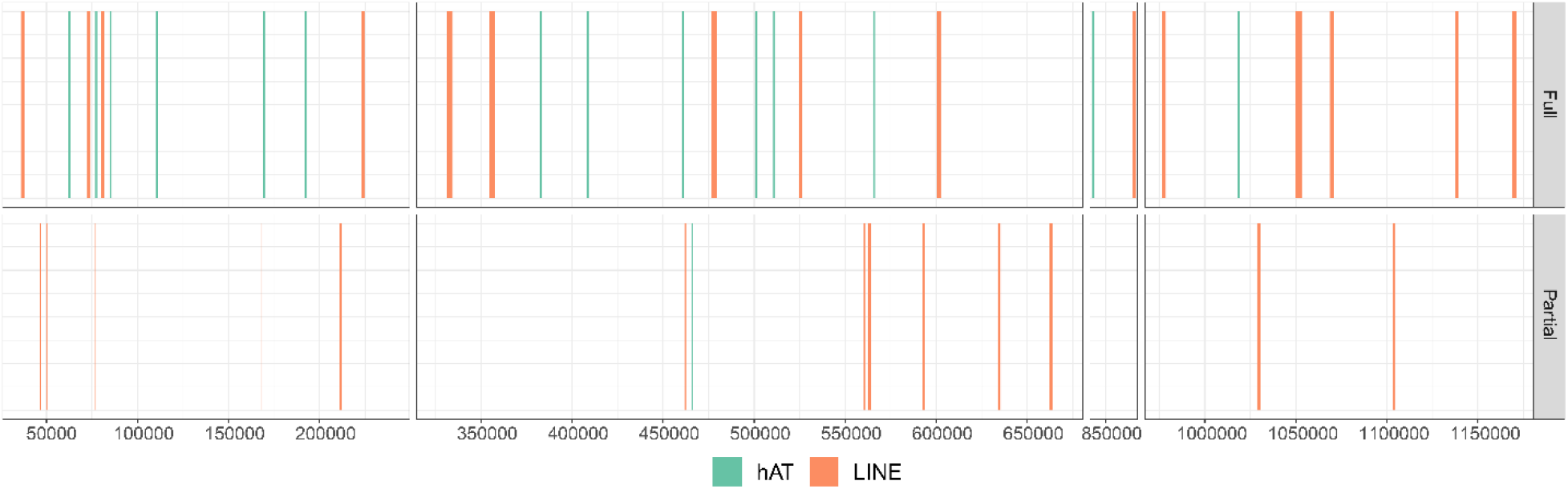
Transposable element insertions in three windows across the genome, separated by full and partial insertions, colored by TE order. Only regions of the genome containing transposable elements are shown, to avoid plotting large gapped regions.

**Supplementary Figure 3:**
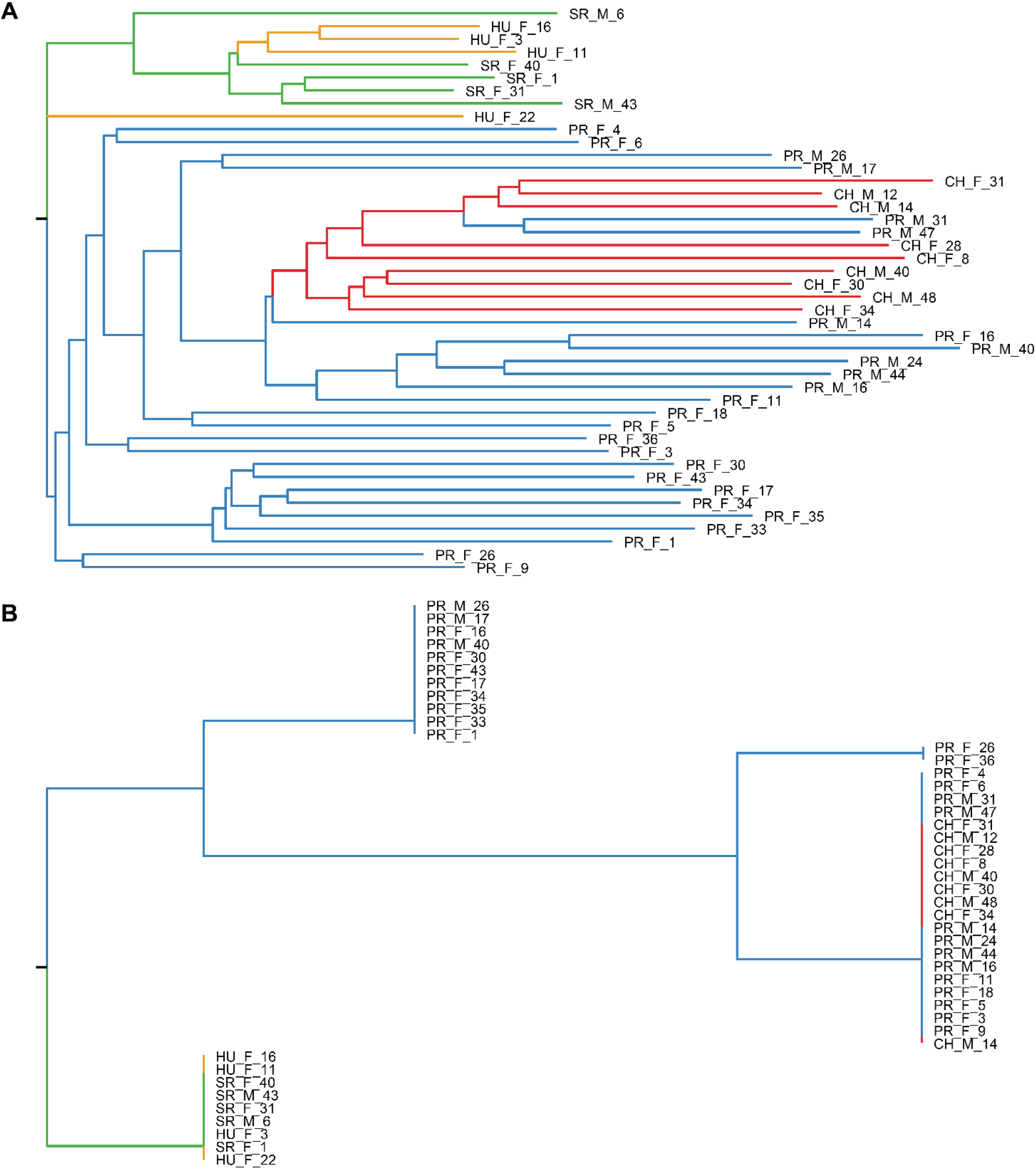
Maximum-Likelihood phylogenies of **A.** Non-prophage *Wolbachia* genes. **B.** the *D. innubila* mitochondria. Branches are colored by the population the tip sample belongs to. Nodes connected to different colored branches are colored by the branch with the most tips.

**Supplementary Figure 4:**
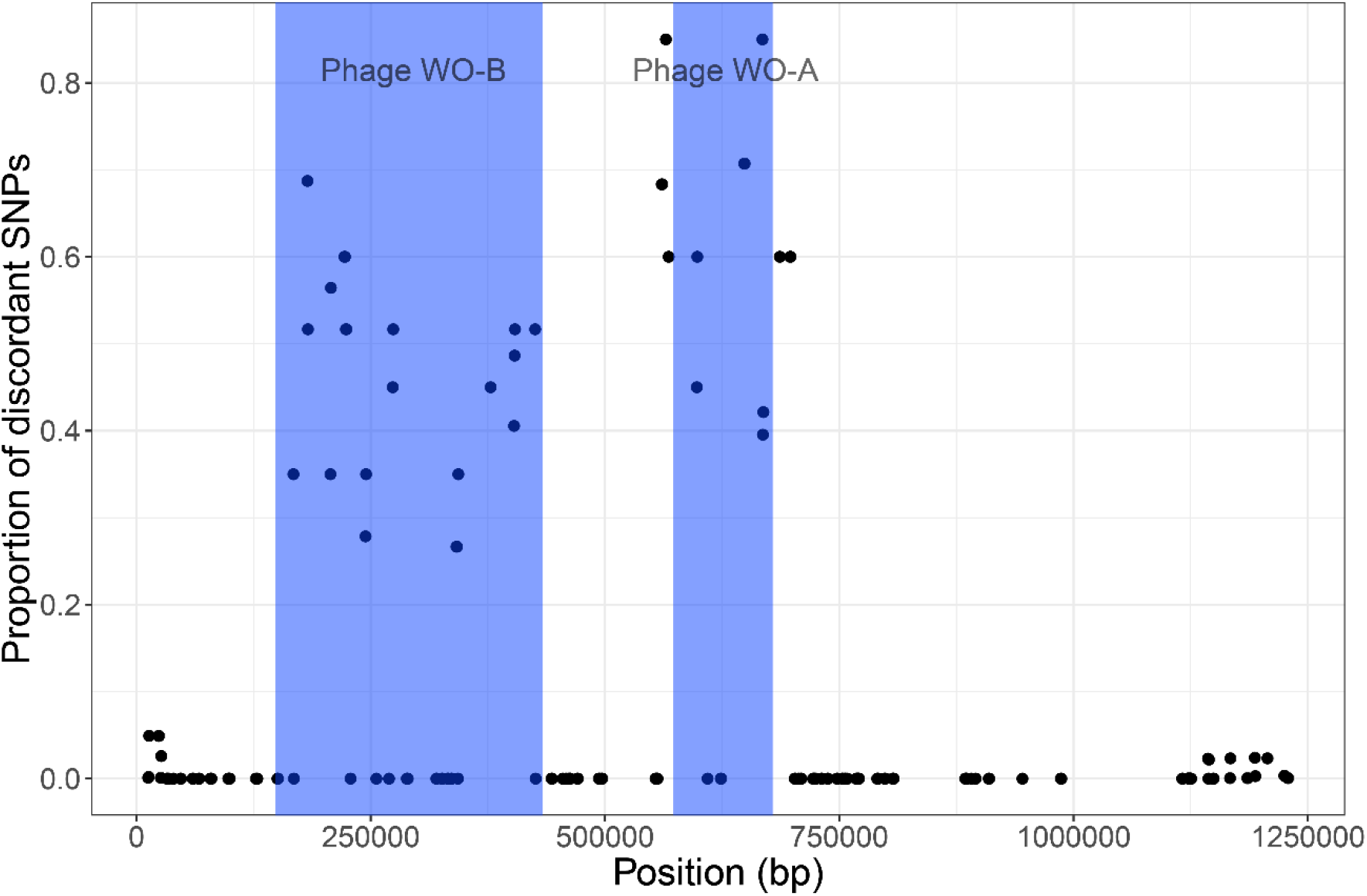
Proportion of SNPs in windows (10kbp windows, sliding 2kbp) across the *w*Inn genome that show discordant inheritance compared to mitochondrial SNPs. Blue shaded blocks highlight the start and end of phage WO portions of the genome. For simplicity we have also shaded the regions between phage blocks of the same type (Figure 1).

